# Dancing Molecules: Rewiring Cooperative Communications within 14-3-3 ζ Docking Proteins

**DOI:** 10.1101/683466

**Authors:** Leroy K. Davis

**Affiliations:** Gene Evolution Project, LLC.; Prairie View A&M University, Cooperative Agriculture Research Center, 700 University Drive, Prairie View, Texas 77446-0518

**Keywords:** Protein Engineering, Synthetic Evolution, Allosteric Interactions

## Abstract

Allosteric engineering may play a key role in novel drug discovery. As allosteric interactions are often associated with disease states where protein active sites are rendered constitutively active. Due to their role in regulating signal transduction in cells, we attempted to rewire cooperative communications within the 14-3-3 ζ docking protein. To avoid disruption of evolutionarily tuned interaction networks, we attempted to do so by applying the “*Fundamental Theory of the Evolution Force*: *FTEF*”. Whereby, we reversed the motion vector of the 14-3-3 ζ C’ terminal tail. We were also able to modify low frequency vibrational modes across 14-3-3 ζ conformational ensembles. Notably, synthetic evolution by *FTEF* anticipated evolution of the 14-3-3 ζ docking protein. And accentuated nonrandom patterns of deformation waves resulting in a gain of function mutation characterized by increased protein flexibility. As well as allowed us to discover a genome encoded spatial arrangement of strain that promotes translation of vibrational motions through protein structural layers. We also discovered a 14-3-3 ζ evolution blueprint that predetermines evolutional fate of the docking protein. The aforementioned suggests that the evolutionary fate of cells may be encoded in the genome, thusly may be predetermined.

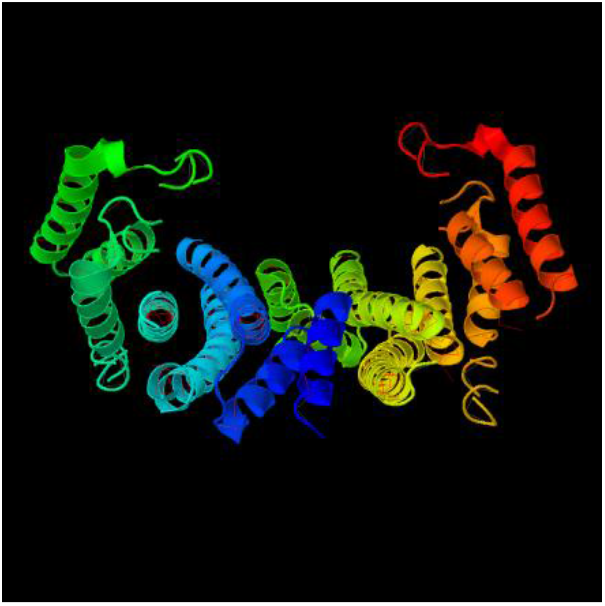

**Significance:** Allostery plays a key role in disease. Wherein, mutations redistribute conformational ensembles and render proteins constitutively active. 14-3-3 docking proteins are key signaling proteins involved in multiple signal pathways that effect cell proliferation, cell cycle and apoptosis. Saliently, the 14-3-3 ζ isoform is associated with neurodegenerative disease, cancer and glaucoma. Thusly, ability to rewire cooperative communications within 14-3-3 ζ offers opportunity for novel drug discovery.

---

Allosteric interactions allow cooperative communication between protein domains by binding of ligands at distant cites leading to altered activity and geometry of their active sites. Allostery can effectively govern signal transduction as proteins comprise of a range of conformational ensembles that allow them to process signals and regulate binding affinity. ^[1]^ The conformer that is most favorable for binding is the one selected. ^[2, 3]^ Allostery arises from events such as binding of ions, drugs and from covalent events such as point mutations. ^[4]^ Where perturbation at any site in the structure leads to shifts in occupancy states across the entire population of preexisting ensembles. ^[3,5,6]^ Conformational ensembles occur as a function of free energy wells with low barriers separating minima wells ^[4]^ that enable molecules to interconvert between conformers. Concomitant free energy landscapes are a function of the spatial arrangement of molecules in three-dimensional space. Whereby, it has been previously suggested that allostery occurs as a function of protein folding. As the protein core promotes a range of ensembles governed by specific packing interactions. ^7]^ Thusly, any perturbation from mutation within the protein’s core would involve reorganization of the evolutionarily tuned interaction network. ^[7]^ Results of mutations are manifested in disease states, where ensembles are redistributed to favor the “ON” state rendering the protein constitutively active. ^[4]^

14-3-3 docking proteins are key regulators that have been reported to interact with over 300 protein binding partners and that effect multiple signaling pathways. ^[43]^ Including pathways associated with cell proliferation, cell cycle and apoptosis. ^[8]^ The family includes (β, ɛ, η, γ, τ, σ, ζ) isoforms. The focus of the current study is rewiring of allosteric interactions in 14-3-3 ζ. The docking protein binds ligands in both its monomeric and dimeric form as dictated by the binding equilibrium. ^[9]^ Binding occurs with protein partners that are in the phosphorylated or unphosphorylated state. As characterized by interaction with serotonin N-acetyl transferase (AANAT) at consensus motif [RRHTLP] via a phosphoserine. As well as binding of raf-1, bad and cdc25 at consensus motif RS*X*pS*X*P. Contrarily, the R18 peptide binds in the unphosphorylated state at consensus sequence WLDL. ^[10, 11]^ Protein and ligand interactions occur within the 14-3-3 ζ amphipathic groove. A highly conserved region within the protein core that is characterized by spatially opposed hydrophobic and polar regions of residues. Mutations in this region have been shown to disrupt ligand interaction. ^[11]^ Due to its many binding partners as well as multiple binding sites it is apparent that 14-3-3 ζ consists of a highly complex interaction of conformational ensembles. Whereby, aberrant mutation may result in abrupt reorganization of the interaction network. ^[7]^ The ability to rewire cooperative communications in 14-3-3 ζ without disrupting natively evolved ensembles would allow altering of cellular pathways. This is significant as 14-3-3 ζ has been associated with neurodegenerative disease and cancer ^[12]^ as well as diseases such as glaucoma via the RhoA signaling pathway ^[43]^. Notably, the ability to engineer allosteric interactions within 14-3-3 ζ docking proteins would offer the potential for novel drug discovery.

The Fundamental Theory of the evolution Force (*FTEF*) is a synthetic evolution approach that allows genes to be written from scratch by identifying genomic building blocks (GBBs) formed over an orthologue/paralogue sequence space during evolution of a gene. GBBs are short sequences that form as evolution artifacts. Whereby, they are identified by evolution force associated with their formation. The evolution force is a compulsion acting at the matter-energy interface that drives molecular diversity and simultaneously promotes conservation of structure and function. ^[13, 14]^ Effects of the evolution force are manifested at all levels of life and is responsible for such processes as the formation of genes and gene networks. ^[13, 14]^ *FTEF* solves for evolution force as a function of four primary evolution engines, (i) evolution conservation, (ii) wobble, (iii) DNA binding state and (iv) periodicity. ^[13, 14]^ When applying *FTEF,* evolution is simulated by assembling genes according to the domain Lego principle. ^[15, 16]^ Whereby, gene sequences are transformed into time based DNA codes according to protein hierarchical structural levels. ^[13, 14]^ Natural selection is simulated by comparing protein structural fingerprints as well as by limiting selection based upon mutation frequency. Gibb’s free energy barriers are also utilized as a selection mechanism, limiting selection to energetically favorable DNA crossovers.

Synthetic evolution by *FTEF* allowed us to engineer 14-3-3 ζ docking proteins that were significantly diverged from a parental *Bos taurus* 14-3-3 ζ docking protein, yet displayed small RMSD variances indicating conservation of 14-3-3 ζ packing interactions. Analysis of allosteric interactions were limited to SYN-AI-1 ζ, the most diverged of the three synthetic proteins engineered in this study. Hydrophobic and coulombic surfaces were conserved within the amphipathic groove located within the protein core. Thusly, synthetic evolution by *FTEF* allowed rewiring of SYN-AI-1 ζ interaction networks without disturbing native conformational ensembles. We analyzed the effect of synthetic evolution on allosteric interactions within the 14-3-3 ζ monomer and homodimer as well as complexes formed with (AANAT). Synthetic evolution by *FTEF* was able to rewire 14-3-3 ζ allosteric interactions and reverse motion vectors by evolving the coulombic surface of the C’ terminal tail. Notably, our theory was also able to identify underlying evolution mechanisms. Whereby, we observed that the genome evolved allosteric interactions within 14-3-3 ζ as a function of spatial arrangement of strain. We also discovered evolution of nonrandom patterns of deformation within the 14-3-3 ζ docking protein and uncovered a genome encoded evolution blueprint that predetermines its evolutional fate.

## 2. Methods

### 2.1 High Performance Computing

Synthetic evolution artificial intelligence (SYN-AI) was performed utilizing the Stampede 2 supercomputer located at the Texas Advanced Computing Center, University of Texas, Austin, Texas. Experiments were performed in the normal mode utilizing SKX compute nodes comprising 48 cores on two sockets with a processor base frequency of 2.10 GHz and a max turbo frequency of 3.70 GHz. Each SKX node comprises 192 GB RAM at 2.67 GHz with 32 KB L1 data cache per core, 1 MB L2 per core and 33 MB L3 per socket. Each socket can cache up to 57 MB with local storage of 144 /tmp partition on a 200 GB SSD.

### 2.2 Simulating DNA Crossovers

SYN-AI simulated evolution by partitioning the parental *Bos taurus* 14-3-3 ζ gene into a DNA secondary code (DSEC) and performing 1 × 10^9^ DNA crossovers within genomic alphabets comprising the (DSEC). DNA hybridizations were performed at 19, 20 and 21 base pairs. Capturing mutations occurring in three open reading frames. DNA hybridization partners were randomly selected across an orthologue/paralogue sequence space constructed by an automated NCBI-Blast. The sequence space comprised of 2.5 × 10^6^ bp of genetic material and of genes at a homology threshold of > 80 percent identity to parental *Bos taurus* 14-3-3 ζ. DNA hybridizations were performed in a buffering solution of 3 mM Mg^2+^ and 1.2 mM dNTP at 328.15° kelvin. ^[17]^ Gibb’s free energy was calculated according to (Owczarzy, 2002) and a penalty assessed for DNA base pair mismatches. ^[18]^

### 2.3 Simulating Natural Selection

Selection was limited to thermodynamically favored DNA crossovers utilizing an inverse tangent sigmoidal function to scale Gibb’s free energy vectors. Free energy vectors were converted to Heaviside nodes by applying an experimental bias and by subsequent transformation utilizing a sinc (x) function in conjunction with a Boolean function. Sequences generating a signal of (1) were considered as genomic building block candidates. A second round of natural selection utilized pattern recognition filters to remove sequences characterized by long stretches of low sequence homology. Thusly, lowering the probability of protein perturbations. A third round of selection limited DNA crossovers to those comprised of evolutionarily favored mutations based upon Blosum 80 mutation frequency. ^[19, 20]^ In a fourth round of selection, DNA crossovers were limited to those displaying evolution earmarks as characterized by (+) molecular wobble vectors. ^[13]^ In a final round of natural selection, DNA crossovers were limited to those characterized by a high magnitude of evolution force.

### 2.4 Engineering Synthetic Super Secondary Structures

Parental super secondary structures were identified utilizing STRIDE knowledge based secondary structure algorithms ^[22]^. Which were utilized to convert the parental 14-3-3 ζ docking protein sequence to a DNA tertiary code (DTER). A round of natural selection was simulated. Where, neural networks selected against sequences that may potentially disrupt protein architecture. Synthetic motifs were engineered by ligation of genomic building blocks randomly selected from genomic alphabet libraries encompassing 5’ to 3’ terminals of parental structures. A cleaving algorithm removed 5’ and 3’ prime overhangs. A second round of natural selection limited selection based on mutation frequency. And a final round of selection imposed a secondary structure homology threshold > 88 percent identity to parental 14-3-3 ζ super secondary structures. A standalone version of PSIPRED 4.0 ^[23]^ was utilized to evaluate protein secondary structure. Synthetic structures passing selection protocols were stored in DTER libraries for writing DNA code.

### 2.5 Writing DNA Code from Scratch

14-3-3 ζ docking genes were written from scratch by walking the DTER. Followed by random selection and ligation of synthetic super secondary structures stored in DTER libraries. SYN-AI constructed a library of 1 × 10^7^ genes that were passed thru a set of neural networks that evaluated closeness of synthetic protein structural states to native states. Wherein, a minimal closeness threshold of > 90 percent identity was set. A subsequent selection limited selection to proteins characterized by naturally occurring mutations based on BLOSUM80 mutation frequency. A further round of natural selection restricted selection to synthetic 14-3-3 docking proteins characterized by mean secondary structure identities located within the top quantile of R quantile quantile normalized vectors. And a final round of selection enriched for functional 14-3-3 ζ docking proteins by comparing closeness of synthetic protein active sites and hydrophobic interfaces to *Bos taurus* 14-3-3 ζ. Wherein, a closeness thresholds of > 90 percent identity was set.

### 2.6 Building Biological Complexes

SYN-AI-1 ζ serotonin N-acetyl transferase biological complexes were built utilizing UCSF Chimera. ^[24]^ Tetrameric and octameric complexes were engineered using Biological Complex I and Biological Complex II PDB crystal structures reported in ^[10]^. Rotamers of native complexes were mutated to those characterizing SYN-AI-1 ζ and C’ terminal residues 215 – 228 of chains A – D deleted. Energy minimization was performed on each chain of synthetic biological complexes utilizing AMBER ^[25 –28]^ with 200 steepest decent steps at a step size of 0.01 Å followed by 20 conjugant gradient steps at 0.01 Å. Hydrogens were added to biological structures utilizing ANTECHAMBER. ^[25, 26]^ Whereby, we applied steric and Gasteiger methods. The process was repeated until good structures were obtained. Structures were solvated utilizing AMBER and by applying Tip3pbox and shell solvent models with a box size of 12.0 Å.

## 3. Results

### 3.1 Rewiring Cooperative Communication in Synthetic 14-3-3 ζ Docking Protein Monomers

According to Reynolds, interaction of ligands at remote sites change the functional site through propagation of subtle conformational changes. ^[28]^ The aforementioned occur through physically contiguous and coevolving amino acids that exist along pre-existing pathways of conformational ensembles. ^[29]^ Herein, we attempt to rewire cooperative communications within the 14-3-3 ζ docking protein by evolving these pre-existing pathways. We performed synthetic evolution by *FTEF* on 14-3-3 ζ monomers, homodimers and biological complexes formed with serotonin N-acetyl transferase (AANAT). Of the three synthetic proteins produced by synthetic evolution, SYN-AI-1 ζ displayed the greatest divergence from parental *Bos taurus* 14-3-3 ζ at 8 percent sequence divergence. The synthetic protein was more closely related to 14-3-3 ζ of *Ophiophagus hannah* and *Anolis carolinensis*. Displaying phylogenetic relationships characterized by score: 281 bits (718), expect: 7e-73, identities: 150/213 (70%) and positives: 165/213 (77%).

The effect of synthetic evolution on allosteric interactions within the SYN-AI-1 ζ monomer was analyzed utilizing the elastic network model (elNemo). Overall 25 normal modes were generated. Three of the observed vibrational motions are illustrated in **Fig. 1**. Where, normal mode 7 was characterized by a “*bend and flex*” motion with the torsos of the monomer flexing closed upon ligand binding. The C’ terminal tail simultaneously thrust forward locking the substrate into the active site. Normal mode 8 was characterized by lateral movement of the C’ terminal tail resulting in an active site transition from an “open” to “closed” state. Normal mode 9 was characterized by a backward thrust and twist of the C’ terminal tail from the forward to resting position. The vibrational vector was inverted from mode 7 and resulted in a reciprocating forward thrust of the C’ terminal tail in transition to the “open” state. All three conformations resulted in unique active site geometries, suggesting they are associated with unique ligands and responsible for regulating different signal transduction pathways.

**Figure 1.**
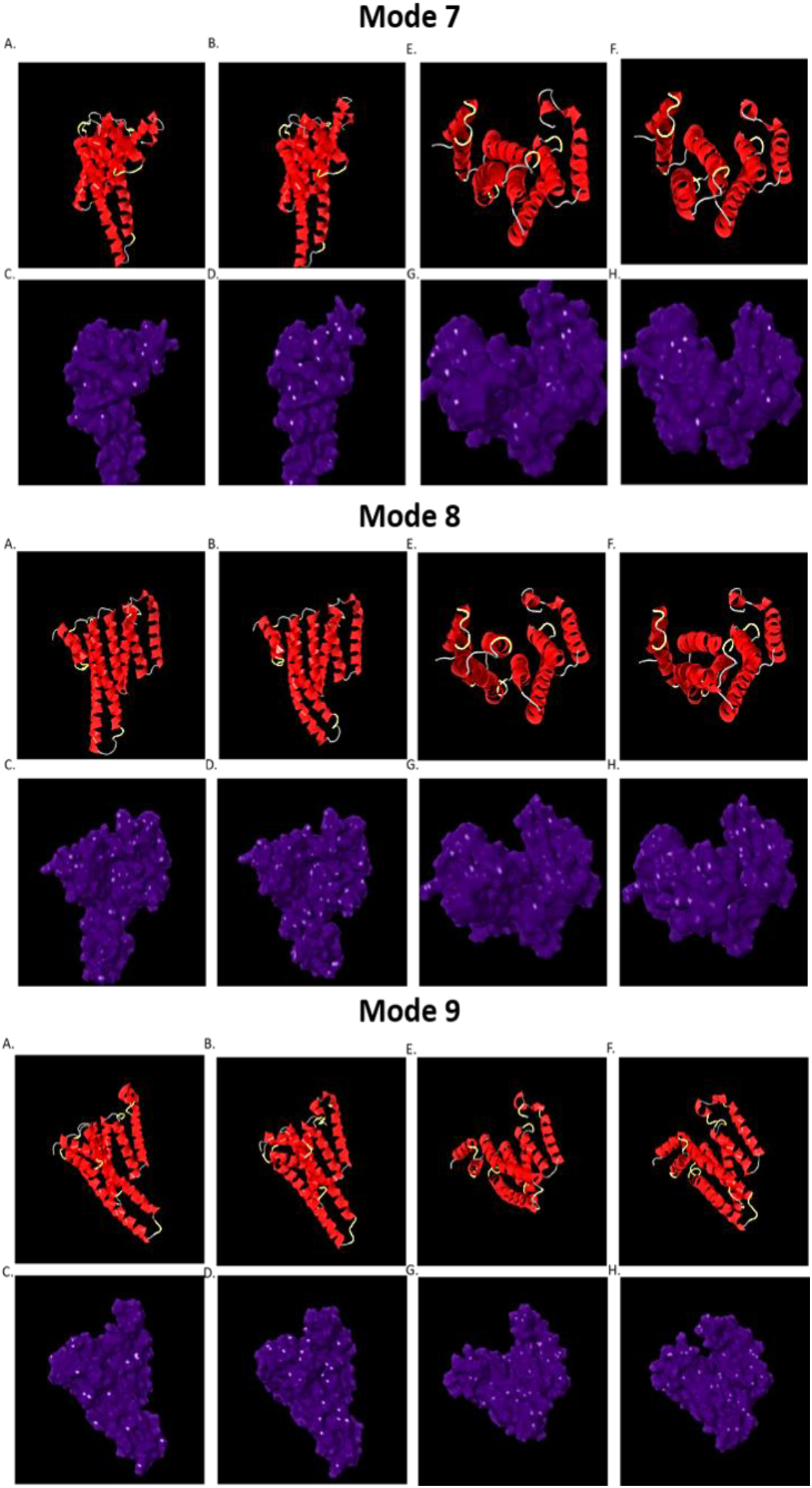
Cooperative Communications in SYN-AI-1 ζ Monomers. Normal mode analysis was performed utilizing the elastic network model, elNemo ^[41, 42]^, with min and max DQ amplitude perturbation of 100 and a DQ step size of 20. Low frequency vibration modes (7 - 9) are depicted above.

### 3.2 The Effects of Synthetic Evolution by FTEF on Cooperative Communication in Synthetic 14-3-3 ζ Homodimers

The human 14-3-3 ζ monomer comprises two residues that are conserved in dimer formation (arginine 21 and lysine 85). These residues are also conserved in *Bos taurus* 14-3-3 ζ. Notably, *FTEF* conserved these residues in all three synthetic proteins. Suggesting that synthetic evolution by *FTEF* did not disrupt 14-3-3 ζ dimerization potential and thereby corroborating COTH ^[38]^ dimerization predictions. Three dimensional SYN-AI-1 ζ homodimer structures were predicted utilizing COTH and normal mode analysis performed utilizing elNemo. Low frequency vibrational motions were conserved in SYN-AI-1 ζ homodimers, **Fig. 2**. Normal mode 7 was translated to the homodimer allowing the molecule to simultaneously bind two ligands and transition from “open” to “closed” conformational states. Signals were transduced by simultaneous forward thrust of juxtaposed C’ terminal tails, **Fig. 2** (A). Normal mode 8 was characterized by SYN-AI-1 ζ monomers displaying vibrational vectors in opposite directions. Where, the right torso of one monomer flexes downward and left torso of the adjacent monomer flexes upward upon ligand binding. Thusly, varying the movement of the C’ terminal tail from Mode 7 and altering active site geometries, **Fig. 2** (B). Normal mode 9 was characterized by simultaneous opening and closing of adjacent monomer active sites. And resulted in altered C’ terminal tail conformations as well as modification of active site geometries from that of modes 7 and 8, **Fig. 2** (C). Our results corroborate that cooperative communication between the SYN-AI-1 ζ active site and C’ terminal tail was not lost in dimeric states.

**Figure 2.**
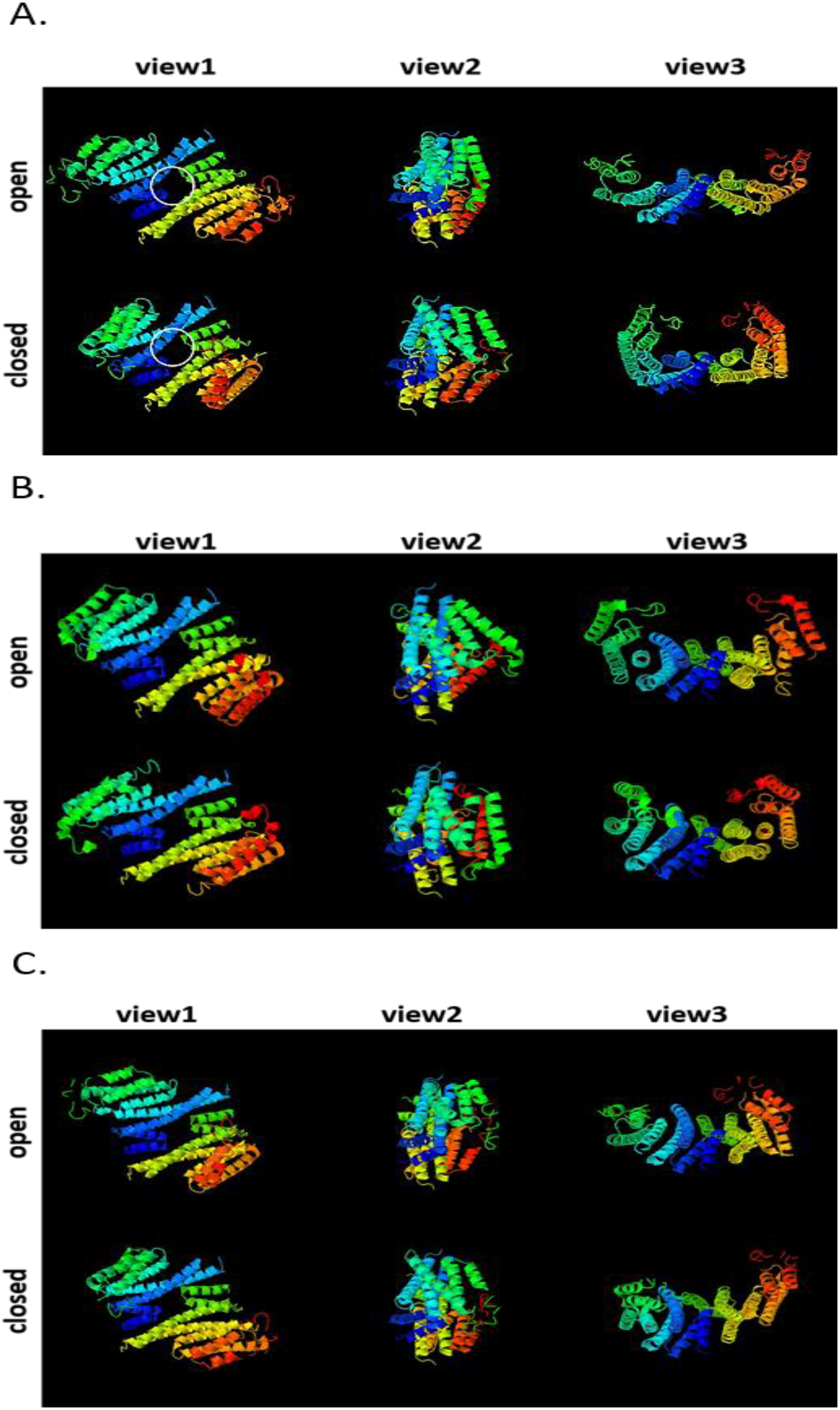
Cooperative communications in SYN-AI-1 ζ homodimers. Structure of synthetic homodimers was predicted utilizing COTH. Normal mode analysis was performed utilizing elNemo with min and max DQ amplitude perturbation of 100 and a DQ step size of 20. Normal mode 7 (**A**), Normal mode 8 (**B**), Normal mode 9 (**C**).

Translation of deformation energy waves through the 14-3-3 ζ structure was analyzed utilizing the Z fluctuation function, (**Eq. 1**). Z fluctuation is the difference in magnitude and net distance fluctuations (**Eq.2** **&** **3**). Thusly, reflects the direction of residue potential energies allowing for characterization of gross potential energy fluctuations within the protein. Where, *L*_*I*_ characterizes the largest fluctuation distance increase and *L*_*D*_ reflects the largest decrease. To optimize newly formed interactions upon ligand binding, atoms move and reorient creating a strain energy that is transduced through protein structural layers. ^[1]^ We analyzed the relationship between14-3-3 ζ deformation vectors (yellow arrows) and strain vectors (red arrows), **Fig.3** (A & B). And utilized two methods to uncover the relationship. Method 1 compared percent C-alpha strain to the percent Z fluctuation. Method 2 compared total C-alpha strain to total Z fluctuation. Method 2 revealed that smaller C-alpha strains translated to grosser energy deformations allowing greater protein flexibility, **Fig. 3** (B). By applying Method 1, we discovered that deformation waves within 14-3-3 ζ form as a function of C-alpha strain vectors that are synonymous to linear potential energy hills, **Fig. 3** (A, C) red arrows. C-alpha strain increases up potential energy hills to a residue of high flexibility followed by a strain inversion (blue/orange asterisk).

**Figure 3.**
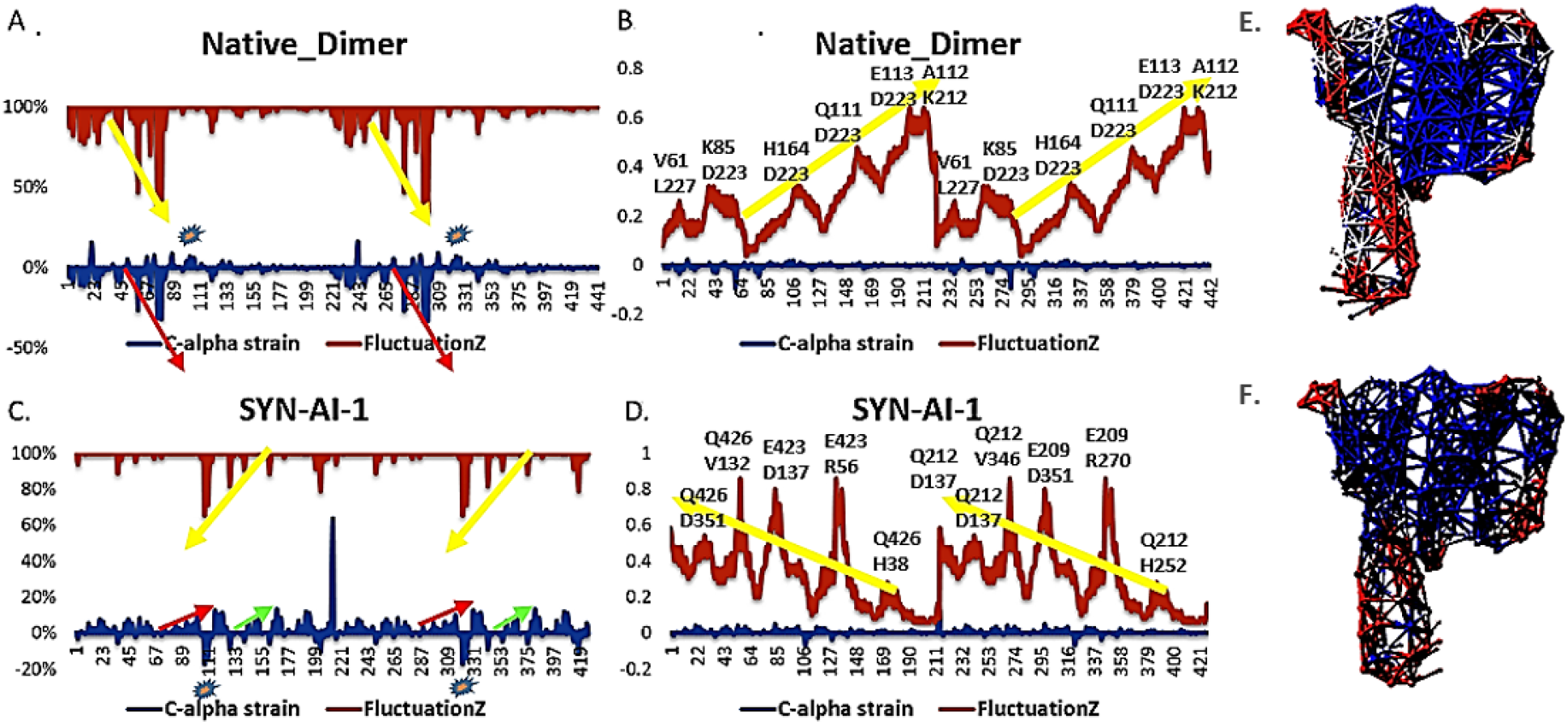
Relationship between Strain and Deformation. The relationship between strain and deformation was analyzed by comparing inter-residue C-alpha strain to Z fluctuation. Normal mode analysis was performed utilizing the elastic network model, elNemo with min and max DQ amplitude perturbation of 100 and a DQ step size of 20. Comparison of percent C-alpha strain to percent Z fluctuation characterizing the parental 14-3-3 ζ homodimer (**A**). Comparison of total C-alpha strain to total Z fluctuation, 14-3-3 ζ homodimer (**B**). Comparison of percent C-alpha strain to percent Z fluctuation, SYN-AI-1 ζ homodimer (**C**). Comparison of C-alpha strain to Z fluctuation, SYN-AI-1 ζ homodimer (**D**). Normal mode analysis was performed utilizing the anisotropic network model, ANM2.1. ^[32]^ Experiments were performed with a distance cutoff between Cα carbons of 15Å and a distance weight factor of 0. Normal mode 7 motion vector of 14-3-3 ζ monomer (**E**). Normal mode vector of SYN-AI-1 ζ monomer (**F**).

Rewiring of SYN-AI-1 ζ cooperative communications was evidenced by reversal of deformation vectors **Fig. 3** (B & D) as well as inversion of potential energy hills. Results were corroborated by the anisotropic network model, **Fig. 3** (E, F).

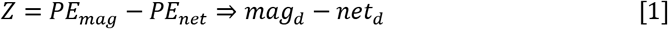

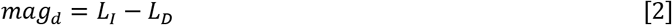

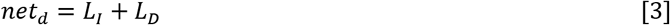

In addition to the occurrence of linear potential energy hills, we discovered additional types of potential energy hills. To avoid complexity, we define linear potential energy hills as Type I and all others as Type II. Rewiring of cooperative communications within SYN-AI-1 ζ is likewise corroborated by inversion and amplification of Type II potential energy hills located between residues 5 – 30, 46 – 67, 113 – 130, 182 – 199 and 210 – 212. Consistent with Type I, they are followed by a strain inversion. Notably, the presence of Type I and II potential energy hills signifies that the genome evolved multiple evolution strategies for transducing deformation waves within 14-3-3 ζ.

### 3.3 Allosteric Interactions in Synthetic 14-3-3 ζ N-acetyl Transferase Biological Complexes

To corroborate rewiring of cooperative communication within SYN-AI-1 ζ biological complexes and transduction of strain via atomic reorientation, we compared inter-residue C-alpha strain to RMSD, **Fig. 4**. Analysis of normal mode 7 indicated an inverse relationship between C-alpha strain and RMSD. Where, large inflections of C-alpha strains located near residues 166 – 170, 395 – 400 and 630 – 640 coincide with RMSD valleys. Further, we superimposed Type I potential energy hills onto the structure. Where, we can visualize the linear increase of strain over helical structures from yellow to red with areas of low strain depicted white. Disordered regions and loops were associated with regions of low strain characterized by strain inversions. Thusly, spatial arrangement of C-alpha strain in biological complexes formed with AANAT was conserved by *FTEF*. Presence of a high flexibility residue at the apex of potential energy hills also occurred in synthetic biological complexes. Our findings support Sinha in that the distribution of conformers is a function of the degree of their molecular flexibility and external conditions in which they reside. ^[4]^

**Figure 4.**
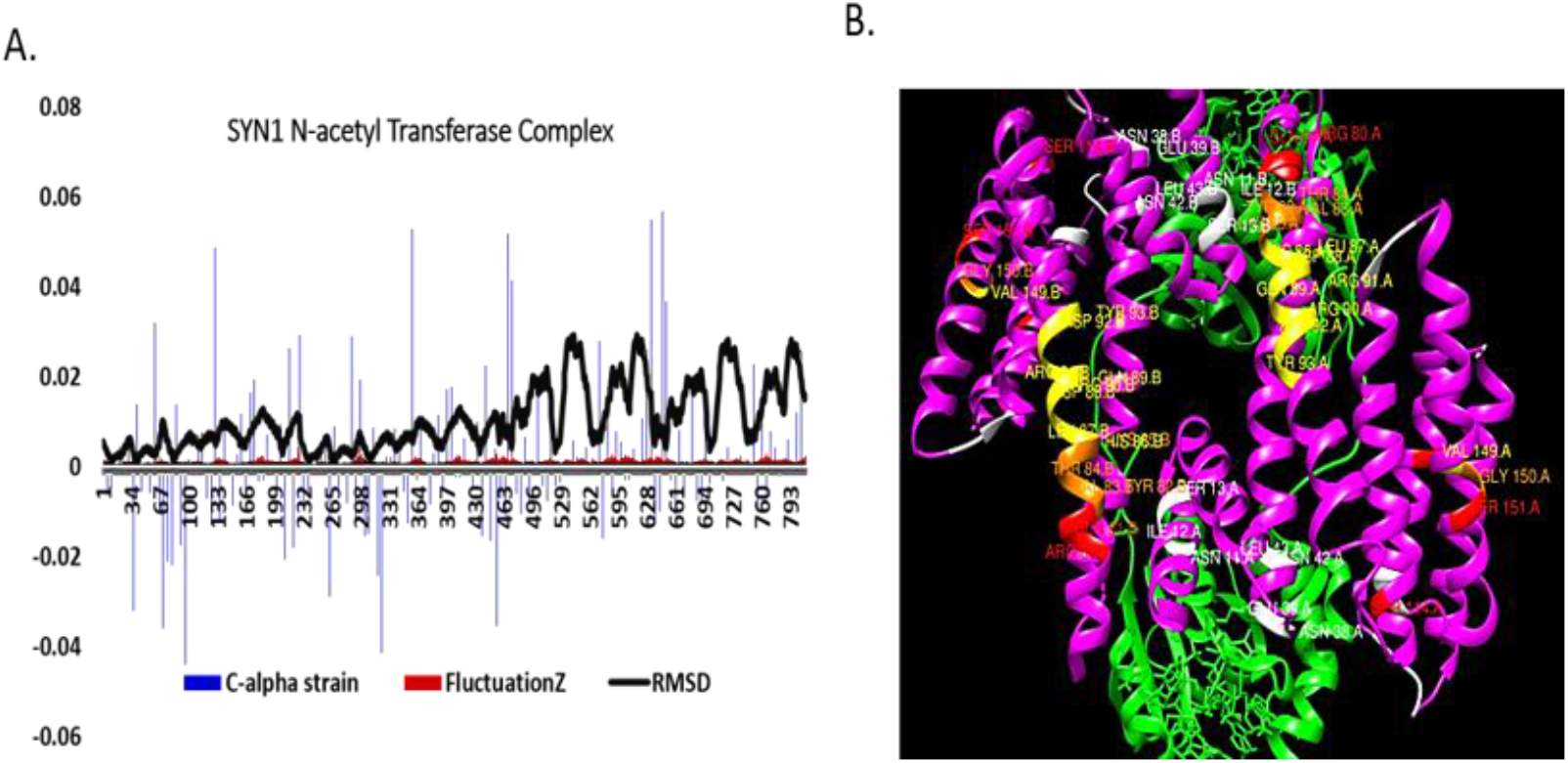
Spatial Arrangement of C-alpha Strain in SYN-AI-1 Biological Complex. Normal mode analysis was performed utilizing elNemo with min and max DQ amplitude perturbation of 100 and a DQ step size of 20. Structure of the SYN-AI-1 ζ serotonin N-acetyl transferase biological complex was predicted utilizing USCF Chimera with energy minimization performed utilizing AMBER. Relationship between strain, deformation and protein packing interactions as characterized by RMSD (**A**). Spatial arrangement of strain in the biological complex (**B**). C-alpha strain increases from yellow to red with areas of low strain depicted white.

Macromolecules consist of ensembles of conformers with a certain distribution which can be described by a free energy landscape. ^[34–36]^ Where, rearrangements between geometrical isomers are considered in terms of transitions between corresponding local minima on the underlying potential energy surface. ^[38]^ Thusly, we corroborated rewiring of 14-3-3 ζ signal transduction by comparing potential energy landscapes of native *Bos taurus* 14-3-3 ζ and SYN-AI-1 ζ homodimers in complex with AANAT. Differences in potential energy landscapes were captured by distance matrices as illustrated in **Fig. 5.** Effects of synthetic evolution are reflected by circled areas. Differences were subtle and location specific and reflect rewiring of allosteric interactions within the SYN-AI-1ζ biological complex. Substantially, there exists no noticeable difference in covariance matrices of native and synthetic biological complexes, **Fig. 6**. Thusly, *FTEF* rewired 14-3-3 ζ cooperative communications without disrupting the global allosteric fingerprint.

**Figure 5.**
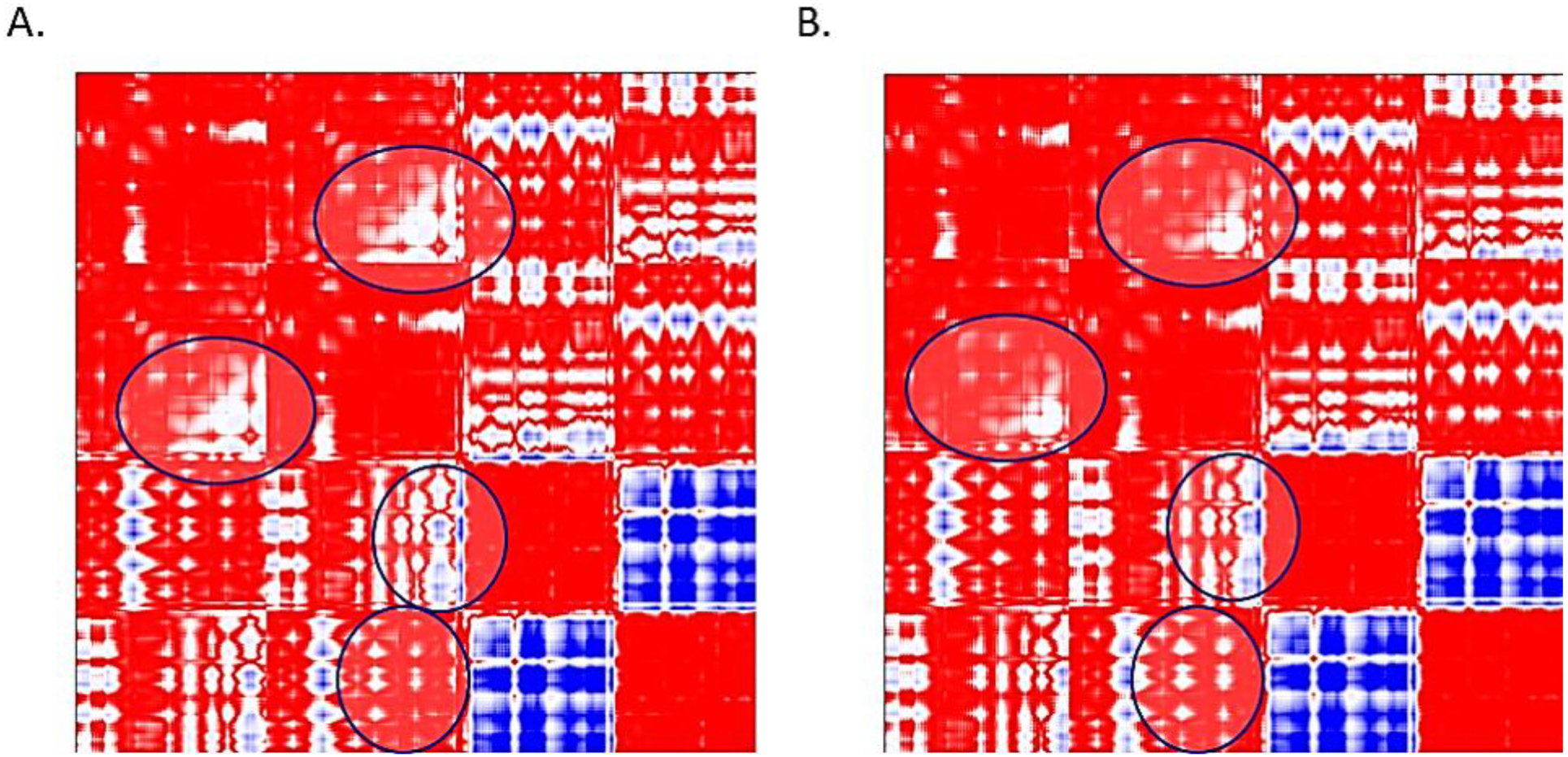
Potential Energy Landscapes. Normal mode analysis was performed utilizing ANM2.1. Experiments were performed with a distance cutoff between Cα carbons of 15 Å and a distance weight factor of 0. The native *Bos taurus* 14-3-3 ζ serotonin N-acetyl transferase biological complex potential energy landscape is characterized by distance matrix (**A**) and the SYN-AI-1 ζ serotonin N-acetyl transferase complex potential energy landscape characterized by distance matrix (**B**).

**Figure 6.**
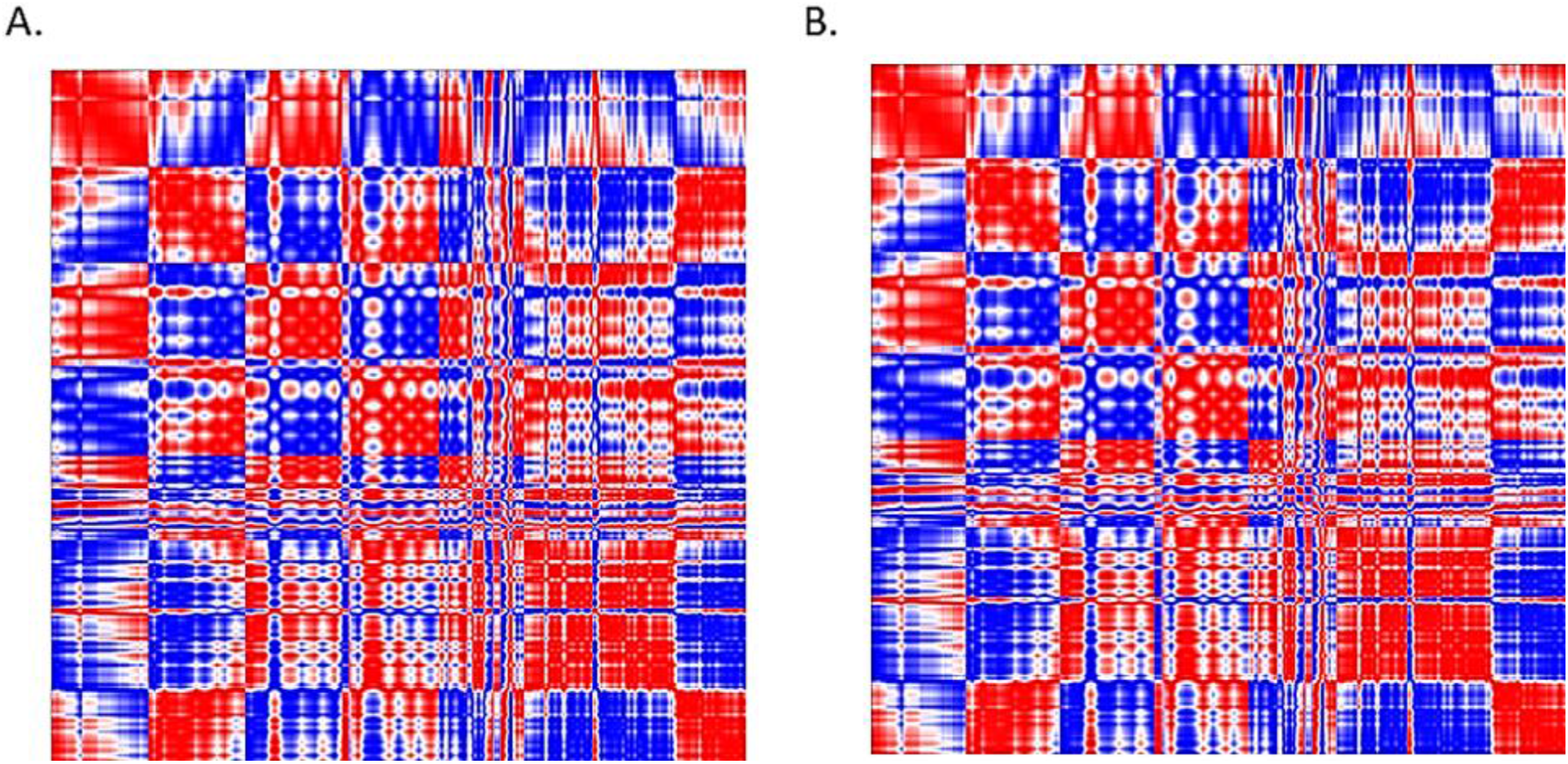
Covariance Matrices. Normal mode analysis was performed utilizing ANM2.1. Experiments were performed with a distance cutoff between Cα carbons of 15 Å and a distance weight factor of 0. The native *Bos taurus* 14-3-3 ζ serotonin N-acetyl transferase biological complex covariance matrix (**A**). SYN-AI-1 ζ serotonin N-acetyl transferase complex covariance matrix (**B)**.

## 4. Discussion

The 14-3-3 ζ docking protein is a key signaling molecule that participates in signal pathways that regulate cell proliferation, cell cycle and apoptosis. Due to the complex interaction of signal pathways and ligand binding events regulated by 14-3-3 ζ, the protein must necessarily comprise of a complex interaction of conformational ensembles that allow for signal integration and ligand differentiation. Formation of conformational ensembles is governed by specific packing interactions that occur within the hydrophobic core. ^[4]^ Whereby, any perturbation from mutation would involve reorganization of evolutionarily tuned interaction networks. ^[7]^ Such mutations occur in disease states resulting in a redistribution of the ensemble to favor the “ON” state rendering the molecule constitutively active. ^[4]^ Due to the role of 14-3-3 ζ in neurodegenerative disease ^[12]^ and cancer ^[39, 40]^, the ability to rewire allosteric interactions without disrupting evolutionarily tuned interaction networks would allow for novel drug discovery.

We attempted to rewire 14-3-3 ζ allosteric interactions utilizing synthetic evolution by *FTEF*. A theory we developed that allows genes to be written from scratch by simulating evolution. ^[13, 14]^ Synthetic evolution by *FTEF* allowed us to engineer three functional 14-3-3 ζ docking proteins. Of which we performed normal mode analysis on SYN-AI-1 ζ the most diverged of the three synthetic proteins. Notably, we were able to rewire cooperative communications within SYN-AI-1 ζ without disrupting native ensembles. Evolution effects varied across conformational ensembles resulting in increased and decreased flexibility in select protein regions, **Figs.** (S1–S4). Suggesting that mutations effect signal pathways differently. Conservation of native ensembles was confirmed by conserved location of deformation peaks across ensembles as illustrated by normal modes (9-12) that were highly conserved, **Fig. S5**. According to Nussinov, biomolecules exist in a range of closely related conformational ensembles. And the conformer most favorable for binding is selected. Our results support Nussinov, as SYN-AI-1 ζ low frequency vibrational motions were characterized by unique transitions to “open” and closed conformational states as well as unique active site geometries.

Majority of SYN-AI-1 ζ low frequency vibrational motions are classified as hinge bending. Suggesting that *FTEF* conserved natively occurring low energy barriers and minima wells. ^[4, 12]^ And that nature engineered these thermodynamic phenomena as a function of C-alpha strain as well as a function of coulombic interactions. It has been postulated that hydrophobic residues located within the protein’s core generate a broad range of conformational isomers. ^[4]^ Notably, *FTEF* conserved 14-3-3 ζ hydrophobic and coulombic surfaces of the amphipathic groove located within the protein’s core, **Fig. S6**. Their conservation allowed rewiring of allosteric interactions without disrupting evolutionarily tuned interaction networks. Contrarily, the coulombic surface of the C’ terminal tail was evolved and resulted in reversal of the Mode 7 motion vector. Suggesting that this region is not as evolutionally conserved as the protein’s core. To grasp the full biological relevance of these allosteric modifications, cellular interconnectedness of the entire network must be considered. ^[1]^ As the effects of allostery are not confined to a single protein but effect the entire signal network, the cell and neighboring cells. ^[29–31]^

During allostery atoms move and reorient creating a strain energy that is transduced through protein structural layers. ^[1]^ Thusly, we compared strain to deformation wave vectors. Our data suggest exploitation of the physical phenomenon of C-alpha strain by the genome to optimize transduction of deformation waves across 14-3-3 ζ structural layers, **Fig. 3**. Whereby, C-alpha strain is associated with formation of potential energy hills. We observed two classes of potential energy hills, linear Type I and nonlinear Type II. The presence of multiple types of potential hills signifies the genome incorporates multiple strategies for transducing deformation within 14-3-3 ζ docking proteins. Their inversion and amplification in SYN-AI-1 ζ corroborates rewiring of allosteric interactions. In both the native and synthetic homodimer, deformations occurred about residues (V, H, D, E, Q, L and R) with the exception of lysine. Suggesting strong evolutional regulation of protein flexibility. Majority of these residues are either polar or charged. Corroborating a critical role of coulombic interactions on cooperative communications. We also identified a correlation between spatial arrangement of C-alpha strain and RMSD, **Fig. 4** (A). The aforementioned suggests evolution of cooperative communications occurs as a function of hydrophobic driven protein folding mechanisms. As RMSD is a reflection of the squeezing out of water molecules during protein compaction and of water density around the core. ^[33]^

To visualize effects of synthetic evolution, we mapped potential energy hills onto the SYN-AI-1 ζ serotonin N-acetyl transferase complex, **Fig. 7**. Type I potential hills were associated with the spine of the docking protein. Allowing deformation waves to transduce down the dorsal spine of SYN-AI-1 ζ as a function of linearly increasing potential energy. Based on strain mappings, the genome designed 14-3-3 ζ docking proteins with a flexible spine that allows monomers rotational freedom during transitioning from “open” to “closed” states. While the upper spine of SYN-AI-1 ζ is very flexible, synthetic evolution by *FTEF* evolved the protein so that the spine’s apex and lower C’ terminal tail (yellow) are inflexible. These mutations allow anchoring of the protein and dorsal spine to prevent distortion during transduction of deformation waves.

**Figure 7.**
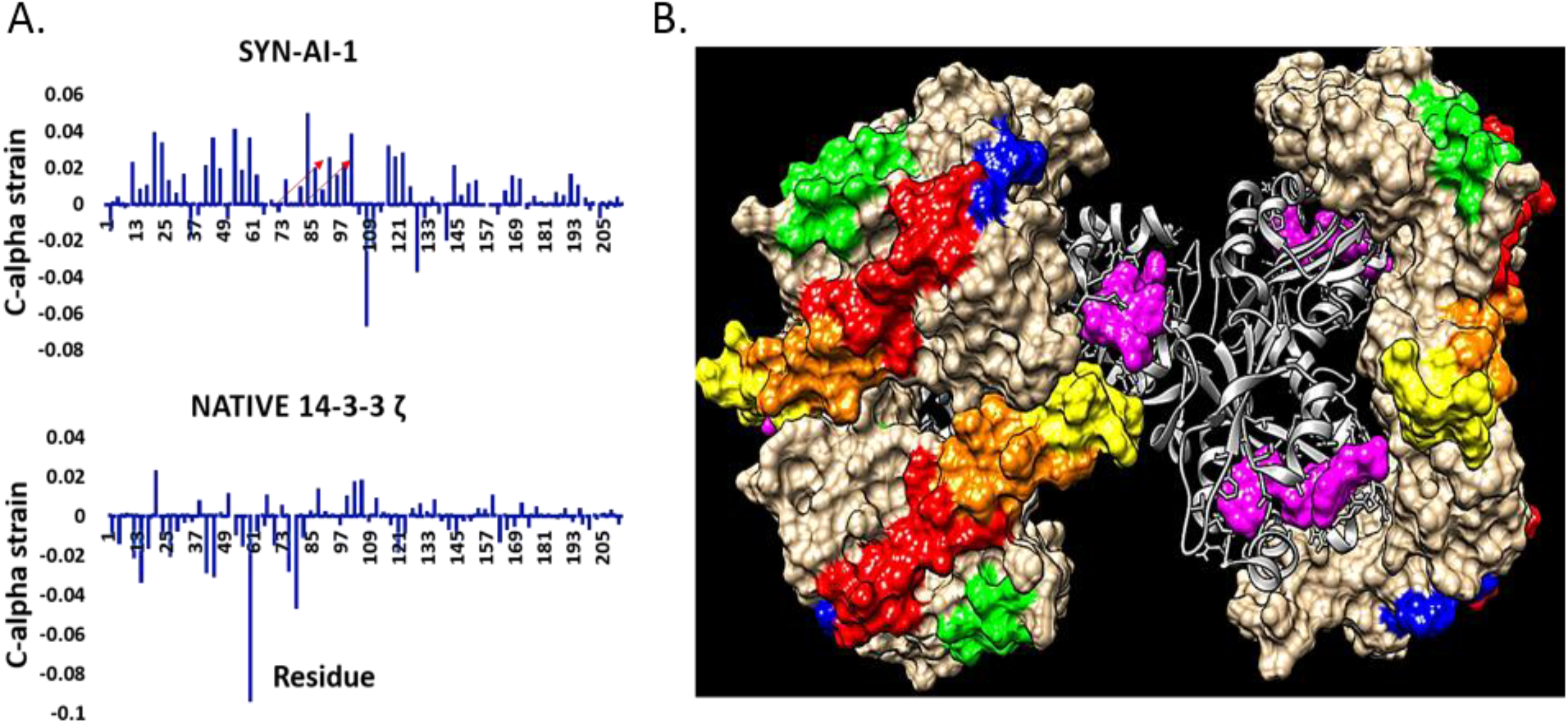
Detection of Genome Encoded Evolution Blueprint. Normal mode analysis was performed utilizing elNemo as previously described. Effects of synthetic evolution were analyzed by comparing C-alpha strain occurring within SYN-AI-1 ζ to that of the parental *Bos taurus* 14-3-3 ζ docking protein (**Left**). The evolution blueprint was visualized by mapping Type I potential energy hills onto the three-dimensional structure of the SYN-AI-1 ζ serotonin N-acetyl transferase biological complex (**Right**). Flexibility increases from yellow to red. Amplified Type I potential hills are depicted green.

Synthetic evolution by *FTEF* displayed anticipatory effects. That resulted in amplification of Type I potential energy hills located between residues 135 – 150 and 255 – 265, **Fig. 3** (green arrows). These potential hills map to a region that allows greater flexibility of the 14-3-3 ζ right torso during binding and locking of the substrate within the active site. Thusly, they are associated with a gain of function mutation. Majority of the residues associated with Type II potential energy hills map to the anterior surface of the SYN-AI-1 ζ spine, **Fig. 8**. Allowing inward compressibility of the C’ terminal tail during hinge bending. Residues associated with Type II potential energy hills also localize adjacent to the spine allowing them to support its movement during “flex and bend” motions. Whereby, we categorized all non-Type I potential energy hills as Type II. It is evident they may be further subcategorized. As there are at least three C-alpha strain mechanisms associated with cooperative motions within synthetic docking protein SYN-AI-1 ζ.

**Figure 8.**
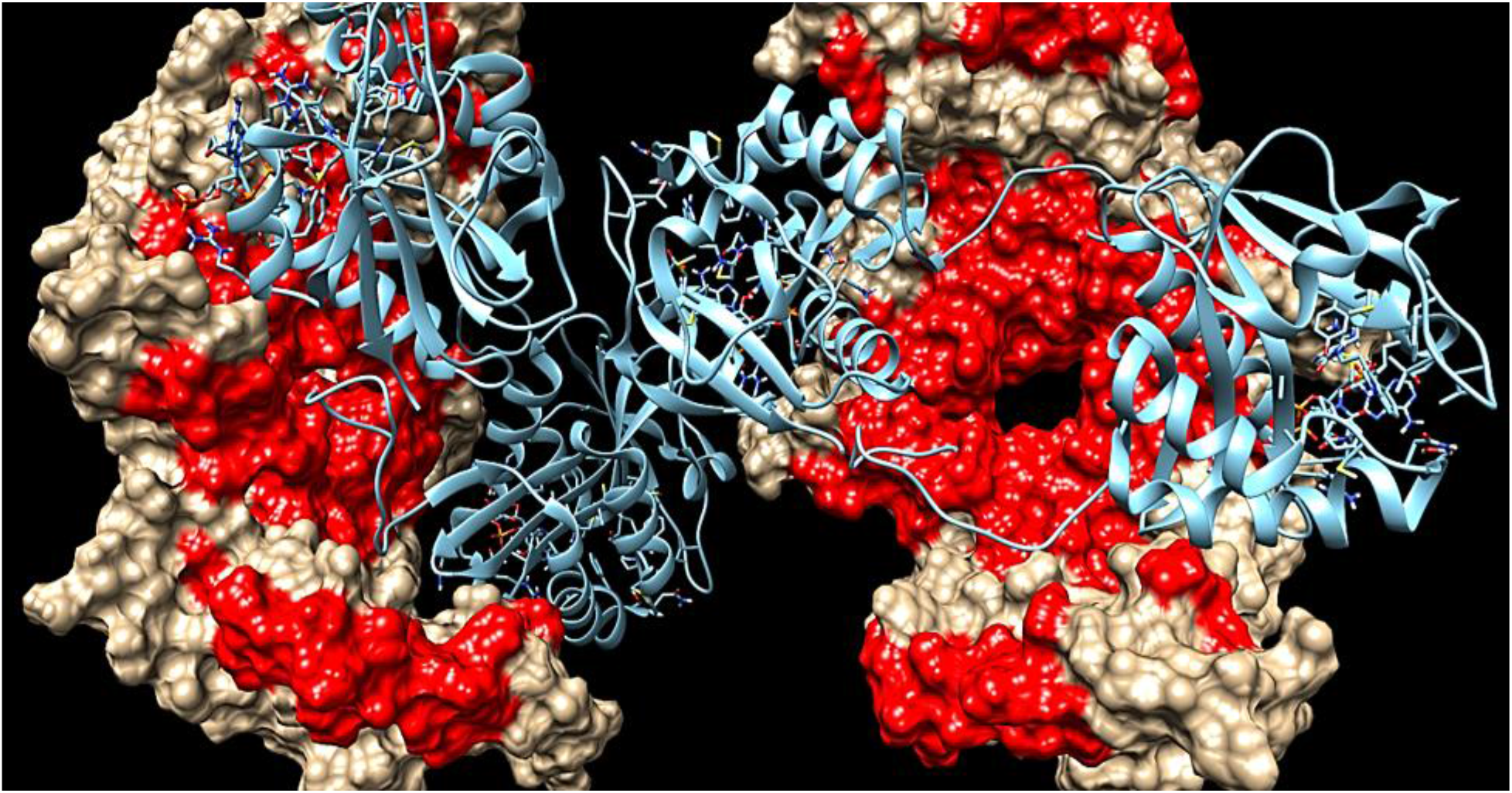
Anticipation of the Evolution Blueprint. Anticipatory effects of synthetic evolution by *FTEF* were analyzed by mapping Type II potential energy hills onto the SYN-AI-1 ζ serotonin N-acetyl transferase complex. Residues associated with Type II potential energy hills are depicted red.

The anticipatory nature of *FTEF* regional selectivity is corroborated by the evolution of nonrandom patterns of C-alpha strain characterizing the SYN-AI-1 ζ homodimer, **Fig. 8** (A, Top). As characterized by the occurrence of two overlapping strain vectors between residues 81 – 100, followed by a strain inversion. The aforementioned form a Type I potential energy hill that maps to the dorsal spine, **Fig. 8** (B) depicted red. Notably, these patterns are not obvious in parental *Bos taurus* 14-3-3 ζ. Our methodology may thusly be a good approach to estimating evolution potential as well as for anticipating evolutional futures of proteins. Synonymous to a builder leaving space for future home expansion, it appears the genome encodes an evolution blueprint containing specifications for gene evolution and diversification as a function of evolutional stringency. Presence of a genome encoded evolution blueprint is further corroborated by observation of nonrandom patterns of evolution within normal mode 14 of SYN-AI-1 ζ, **Fig. S 1**. Where, synthetic evolution by *FTEF* introduced three parallel Type I motion vectors (3, 4, 5), **Fig. S.1** (C), of equal slope and two countercurrent parallel motion vectors (1, 3). These resulted in increased flexibility between residues 13 – 100 characterized by deformation peaks A and B. Whereby, introduction of motion vectors 8 and 9 was responsible for formation of deformation waves C, D and E that also resulted in a gain of function. The continued formation of Type I potential hills during evolution of the 14-3-3 ζ docking protein corroborates the presence of a genome encoded evolution blueprint. Notably, when expounded on, presence of such a blueprint would infer that evolutionary fates of cells may also be predetermined.

To further confirm anticipatory effects of synthetic evolution by *FTEF*, we evaluated C-alpha strain of synthetic proteins SYN-AI-2 ζ and SYN-AI-3 ζ. These proteins are 7 % diverged from parental 14-3-3 ζ compared to SYN-AI-1 ζ that is 8 % diverged from the parental docking protein. Notably, the Type I potential hill located between residue 73 – 110 of SYN-AI-2 ζ is similar to the one observed in SYN-AI-1 ζ. Where, there is a linear increase of flexibility over the SYN-AI-2 ζ dorsal spine. Further supporting evidence of a genome encoded evolution blueprint. Additionally, Type I motion vectors were introduced to different regions of the docking protein. As expected, due to significant levels of divergence characterizing synthetic proteins their allosteric fingerprints vary. Prominently, nonrandom patterns of C-alpha strain occurred in both synthetic docking proteins as depicted by arrows, **Fig. 9**.

**Figure 9.**
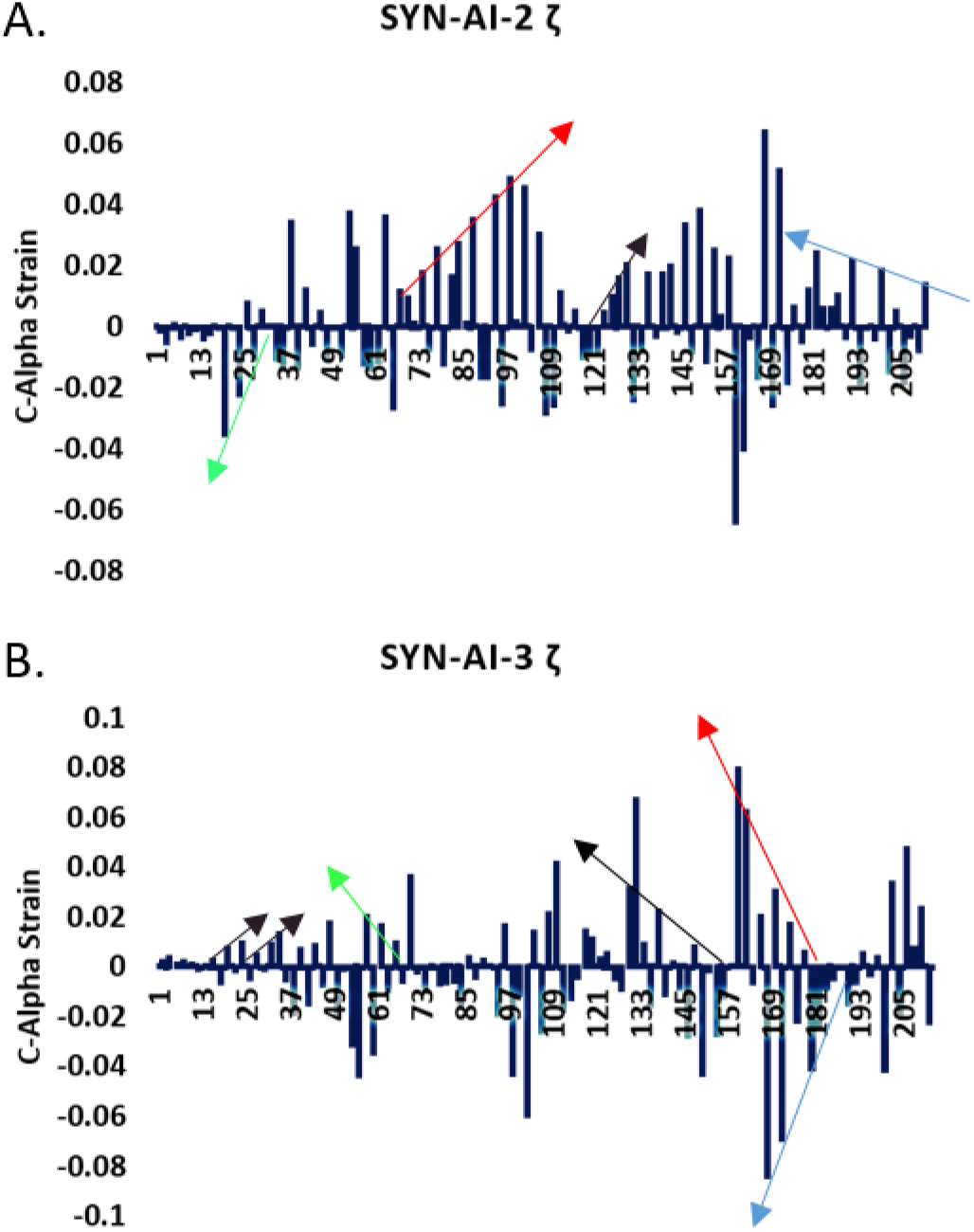
Nonrandom Patterns of Strain. Normal mode analysis was performed utilizing elNemo as previously described. SYN-AI-2 ζ (**Top**). SYN-AI-3 ζ (**Bottom**). Type I strain vectors depicted by arrows.

## 5. Conclusions

By applying the “*Fundamental Theory of the Evolution Force: FTEF*”, we evolved cooperative communications in 14-3-3 ζ without disturbing natively evolved conformational ensembles. As corroborated by conservation of native *Bos taurus* 14-3-3 ζ allosteric fingerprints. Notably, *FTEF* showed the potential to capture intrinsic evolution mechanisms. As *FTEF* provided data that the genome encodes the translation of deformation waves through protein structural layers as a function of spatial arrangement of C-alpha strain. And that strain shares a relationship with RMSD. Thusly, corroborating dependency of cooperative communications on protein folding dynamics that are a function of specific packing interactions. *FTEF* conserved the hydrophobic surface of the 14-3-3 ζ docking protein enabling introduction of mutations without disturbing evolutionarily tuned interaction networks. Wherein, synthetic evolution by *FTEF* resulted in gain of function mutations that increased flexibility of the 14-3-3 ζ dorsal spine and adjacent areas that support the spine during “Hinge Bending” motions. Notably, due to nonrandom patterns of linearly increasing C-alpha strain down the 14-3-3 ζ dorsal spine as a result of synthetic evolution by FTEF, we conclude that the genome encodes an evolution blueprint that allows diversification and expansion of 14-3-3 ζ function. Presence of such a blueprint suggests that evolutionary fates of cells may be encoded in the genome, thusly may be predetermined.

## Acknowledgements

This work used Extreme Science and Engineering Discovery Environment (XSEDE) that is supported by National Science Foundation grant ACI-1548562. The authors acknowledge the Texas Advanced Computing Center (TACC) at the University of Texas at Austin for providing HPC resources. URL: http://www.tacc.utexas.edu

## Supplemental Figures

**Figure S.1.**
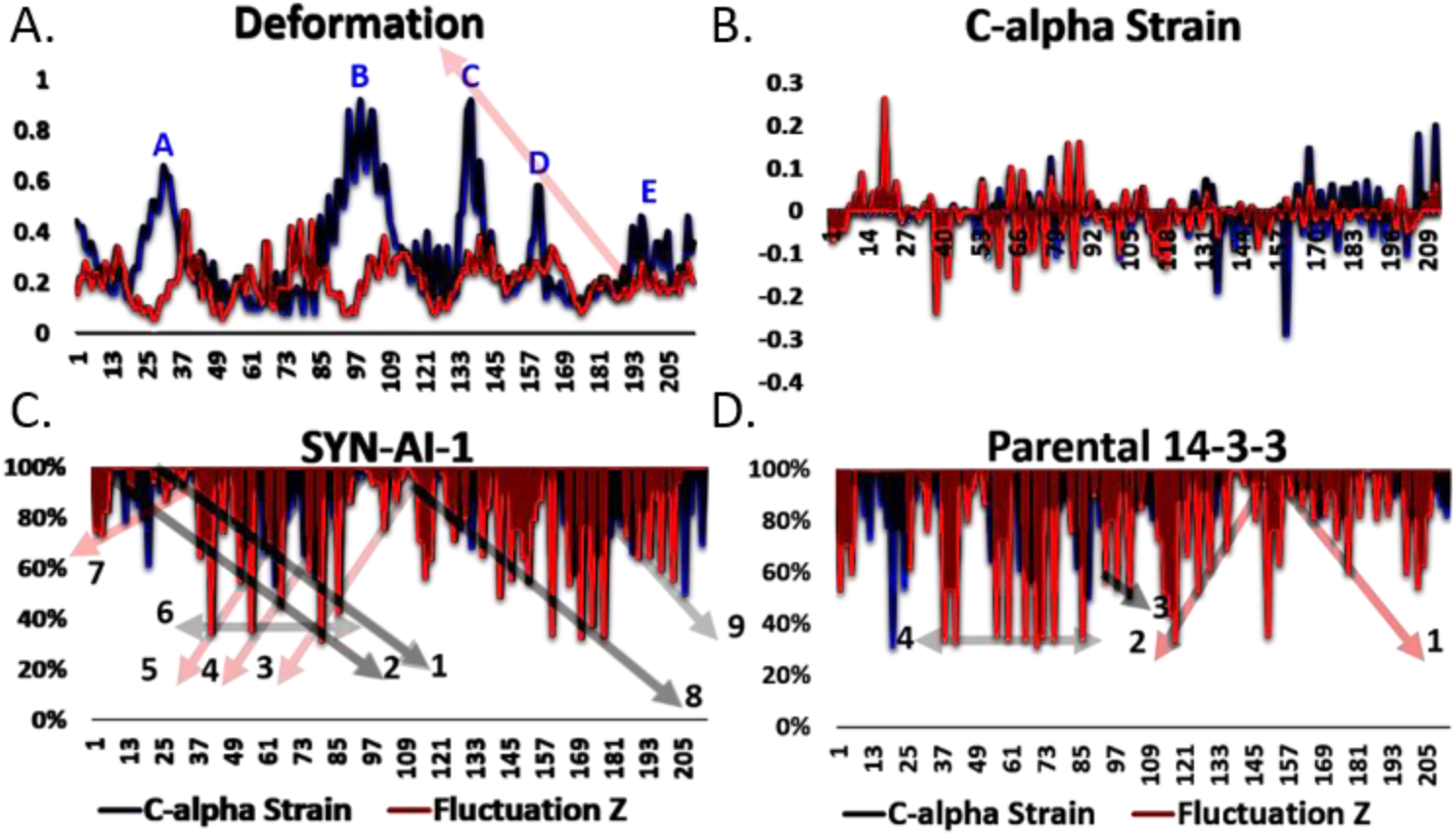
Rewired Allosteric Interactions in Normal Mode 14. Normal mode analysis was performed utilizing elNemo with min and max DQ amplitude perturbation of 100 with a DQ step size of 20. Z fluctuation analysis of deformation vectors, SYN-AI-1 ζ (Blue) and Parental *Bos taurus* 14-3-3 ζ (Red), (**A**). C-alpha strain of SYN-AI-1 ζ (Blue) compared to *Bos taurus* 14-3-3 ζ (Red), (**B**). The relationship between C-alpha strain and deformation was analyzed by comparing percent C-alpha strain to Z fluctuation in SYN-AI-1 ζ (**C**) and *Bos taurus* 14-3-3 ζ (**D**).

**Figure S.2.**
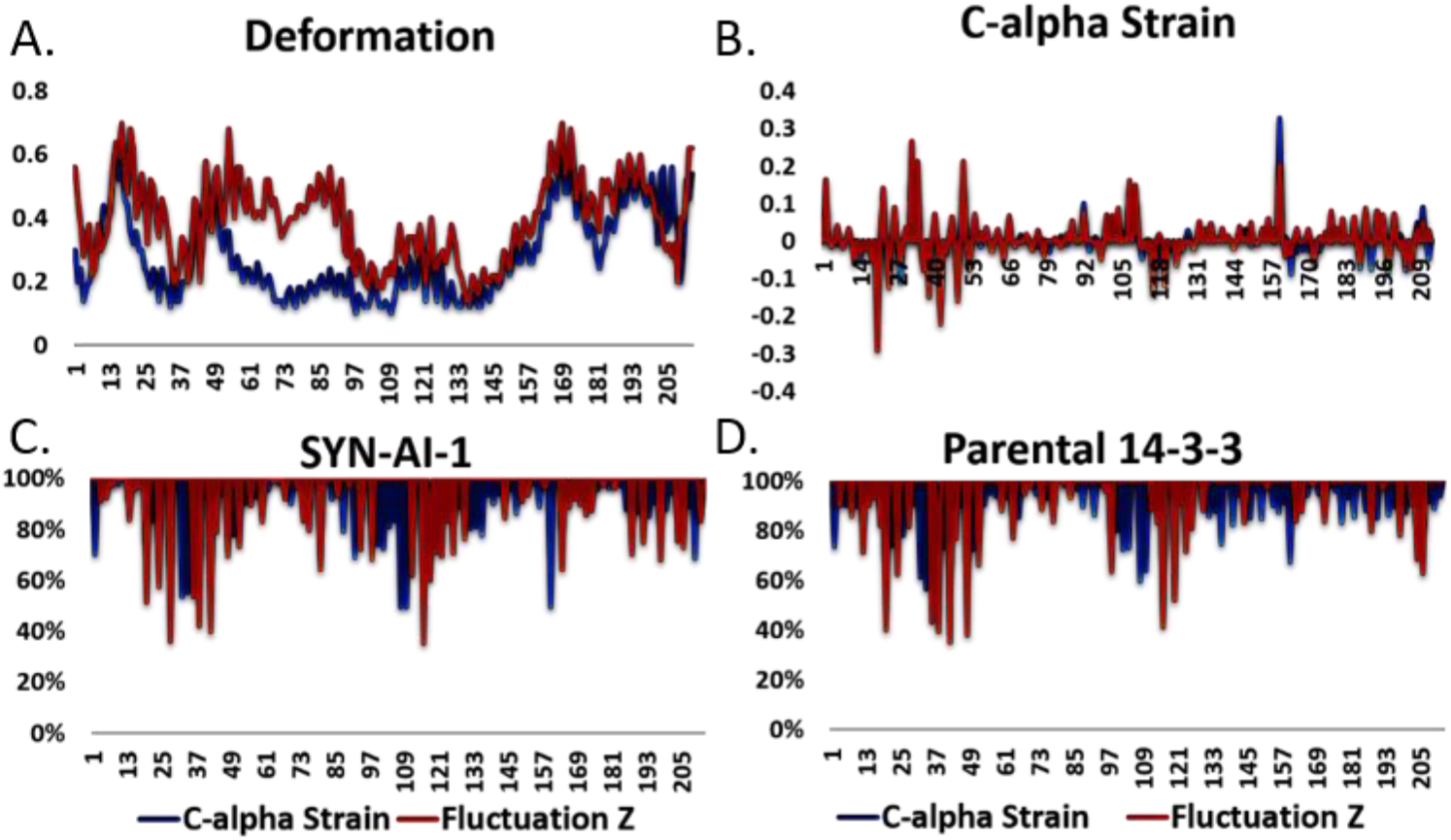
Rewired Allosteric Interactions in Normal Mode 13. Normal mode analysis was performed utilizing elNemo with min and max DQ amplitude perturbation of 100 with a DQ step size of 20. Z fluctuation analysis of deformation vectors, SYN-AI-1 ζ (Blue) and Parental *Bos taurus* 14-3-3 ζ (Red), (**A**). C-alpha strain of SYN-AI-1ζ (Blue) compared to *Bos taurus* 14-3-3 ζ (Red), (**B**). The relationship between C-alpha strain and deformation was analyzed by comparing percent C-alpha strain to Z fluctuation in SYN-AI-1 ζ (**C**) and *Bos taurus* 14-3-3 ζ (**D**).

**Figure S.3.**
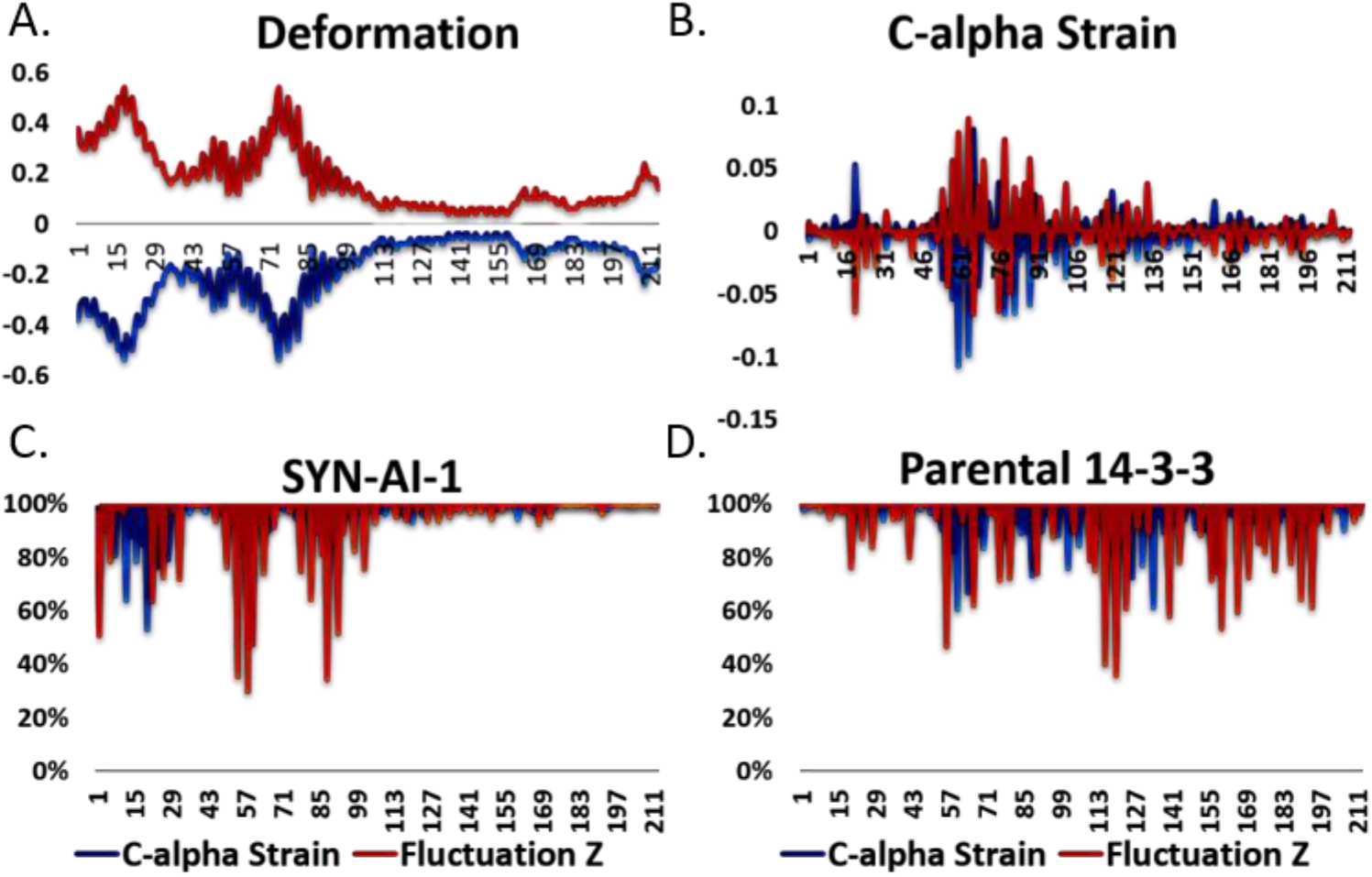
Rewired Allosteric Interactions in Normal Mode 8. Normal mode analysis was performed utilizing elNemo with min and max DQ amplitude perturbation of 100 with a DQ step size of 20. Z fluctuation analysis of deformation vectors, SYN-AI-1 ζ (Blue) and Parental *Bos taurus* 14-3-3 ζ (Red), (**A**). C-alpha strain of SYN-AI-1 ζ (Blue) compared to *Bos taurus* 14-3-3 ζ (Red), (**B**). The relationship between C-alpha strain and deformation was analyzed by comparing percent C-alpha strain to Z fluctuation in SYN-AI-1 ζ (**C**) and *Bos taurus* 14-3-3 ζ (**D**).

**Figure S.4.**
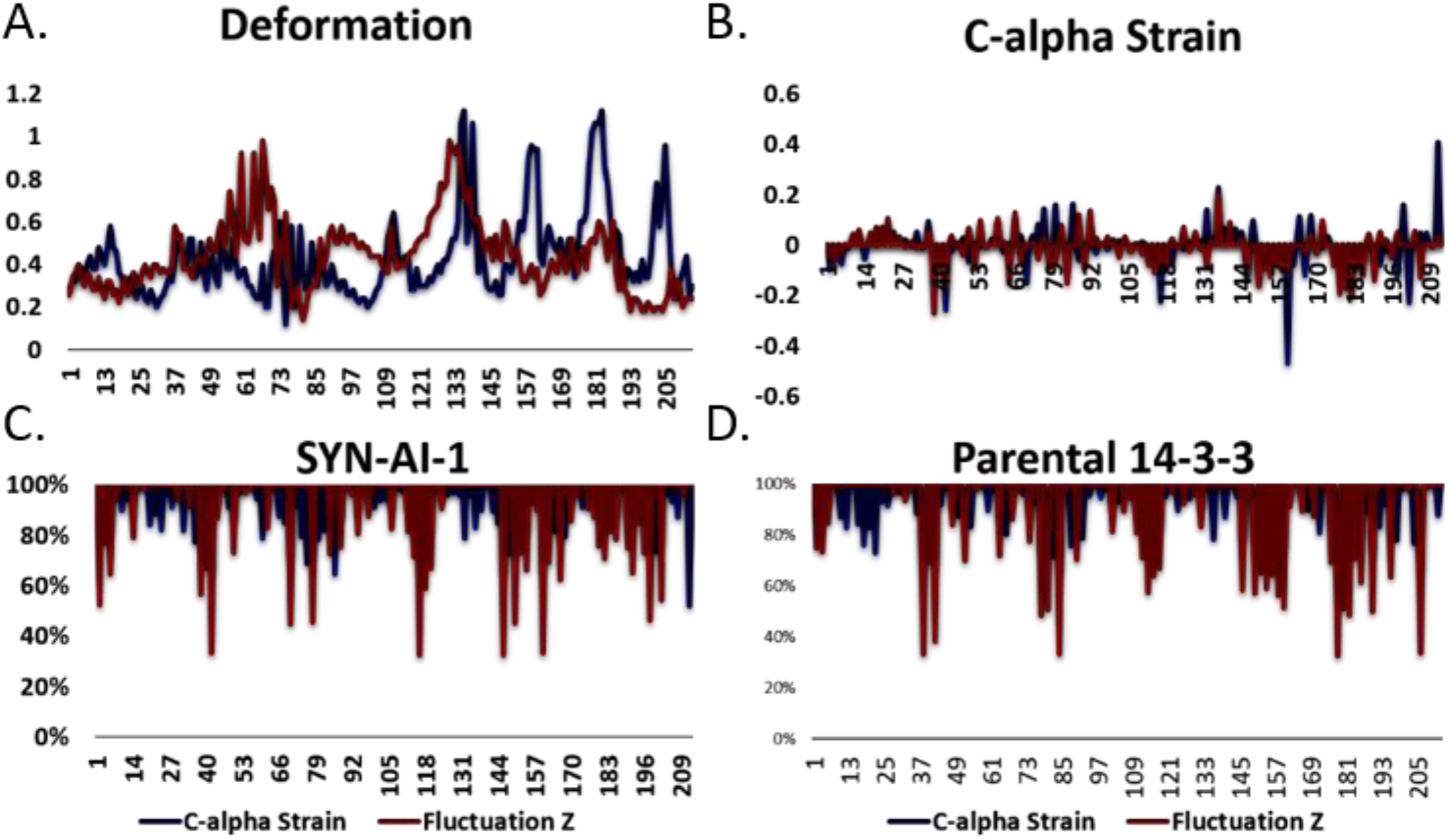
Rewired Allosteric Interactions in Normal Mode 17. Normal mode analysis was performed utilizing elNemo with min and max DQ amplitude perturbation of 100 with a DQ step size of 20. Z fluctuation analysis of deformation vectors, SYN-AI-1 ζ (Blue) and Parental *Bos taurus* 14-3-3 ζ (Red), (**A**). C-alpha strain of SYN-AI-1 (Blue) compared to *Bos taurus* 14-3-3 ζ (Red), (**B**). The relationship between C-alpha strain and deformation was analyzed by comparing percent C-alpha strain to Z fluctuation in SYN-AI-1 ζ (**C**) and *Bos taurus* 14-3-3 ζ (**D**).

**Figure S.5.**
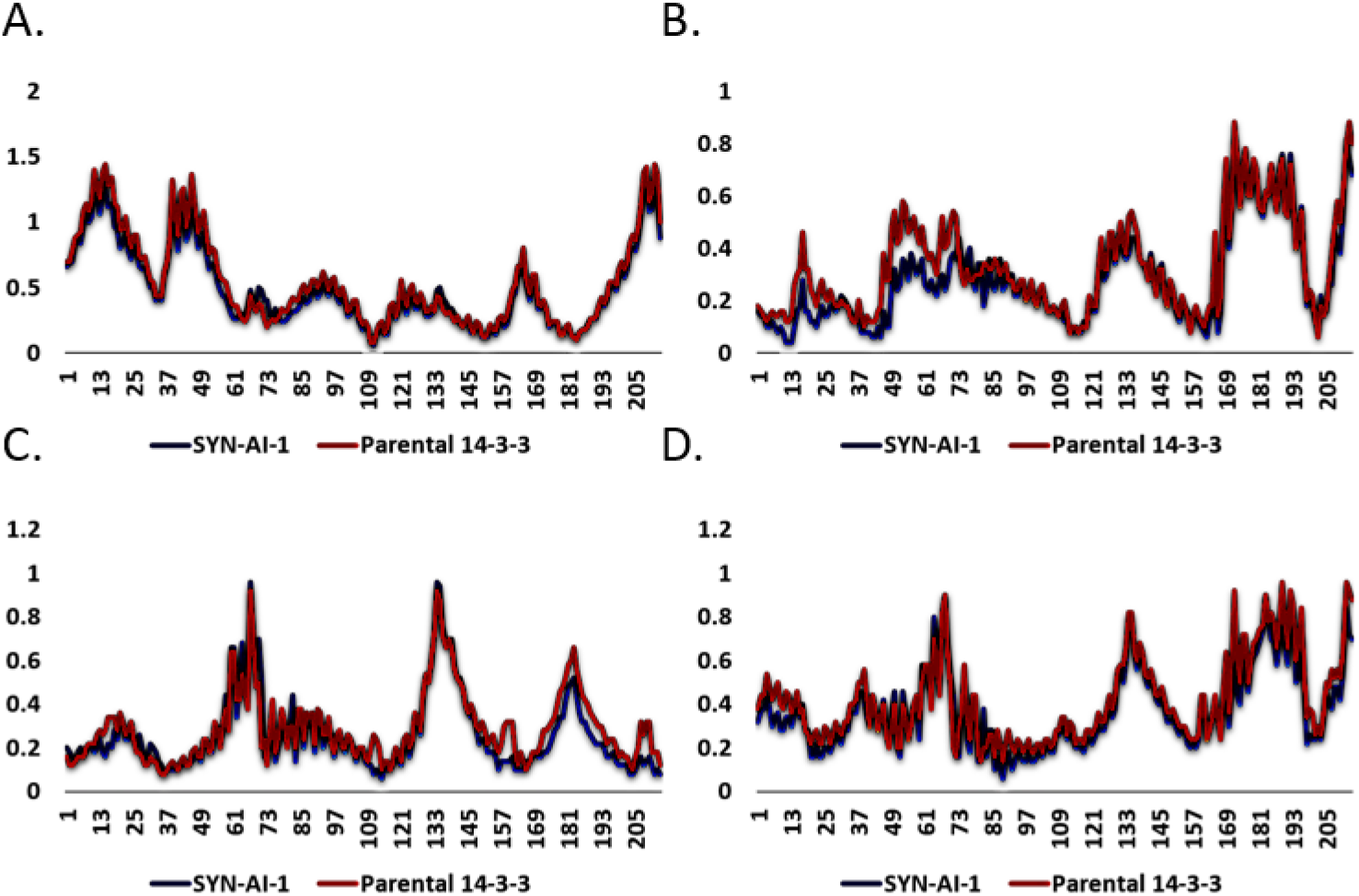
Conservation of Native Conformational Ensembles. Normal mode analysis was performed utilizing elNemo with min and max DQ amplitude perturbation of 100 with a DQ step size of 20. Comparison of native *Bos taurus* 14-3-3 ζ deformation versus synthetic protein SYN-AI-1 ζ. Normal mode 9 (**A**). Normal mode 10 (**B**). Normal mode 11 (**C**). Normal mode 12 (**D**).

**Figure S6.**
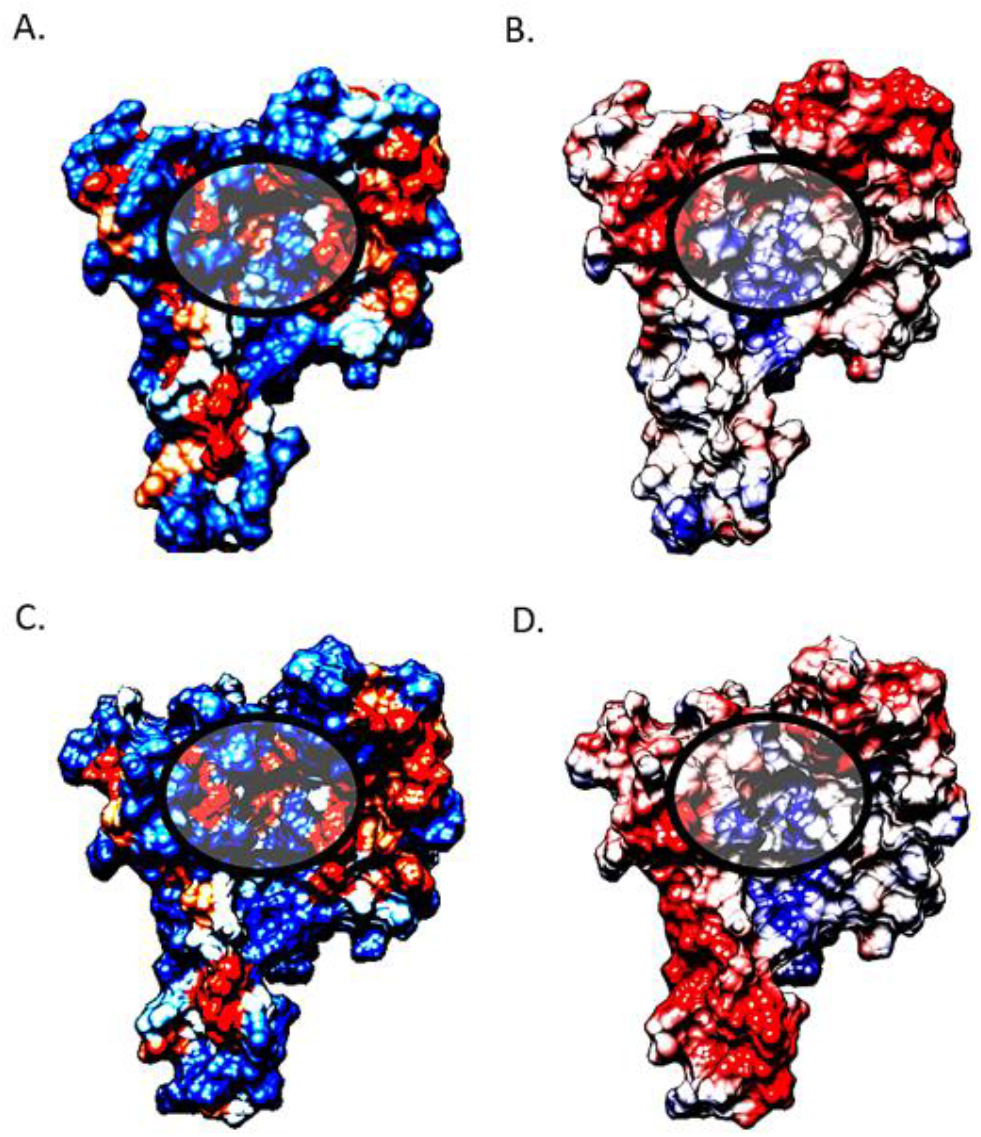
Conservation of 14-3-3 ζ Hydrophobic and Coulombic surfaces. Three-dimensional structure and PDB coordinates of synthetic protein SYN-AI-1 ζ were predicted utilizing I-TASSER, Zhang Laboratory, University of Michigan. Docking protein hydrophobic and coulombic surfaces were analyzed utilizing USCF Chimera. The parental *Bos taurus* 14-3-3 ζ structure was truncated from residues 215 - 228 and compared to the synthetic protein. Hydrophobic surface map of parental *Bos taurus* 14-3-3 ζ (**A**). Coulombic surface of parental 14-3-3 ζ (**B**). Hydrophobic surface map of synthetic protein SYN-AI-1 ζ (**C**). SYN-AI-1 ζ coulombic surface map (**D**). The 14-3-3 ζ amphipathic groove is denoted by a circle.

